# Identification of potential biomarkers associated with pathogenesis of primary prostate cancer based on meta-analysis approaches

**DOI:** 10.1101/2020.03.05.978205

**Authors:** Neda Sepahi, Mehrdad Piran, Mehran Piran, Ali Ghanbariasad

## Abstract

Worldwide prostate cancer (PCa) is recognized as the second most common diagnosed cancer and the fifth leading cause of cancer death among men globally. Rising incidence rates of PCa have been observed over the last few decades. It is necessary to improve prostate cancer detection, diagnosis, treatment and survival. However, there are few reliable biomarkers for early prostate cancer diagnosis and prognosis. In the current study, systems biology method was applied for transcriptomic data analysis to identify potential biomarkers for primary PCa. We firstly identified differentially expressed genes (DEGs) between primary PCa and normal samples. Then the DEGs were mapped in Wikipathways and gene ontology database to conduct functional categories enrichment analysis. 1575 unique DEGs with adjusted p-value < 0.05 were achieved from two sets of DEGs. 132 common DEGs between two sets of DEGs were retrieved. The final DEGs were selected from 60 common upregulated and 72 common downregulated genes between datasets. In conclusion, we demonstrated some potential biomarkers (FOXA1, AGR2, EPCAM, CLDN3, ERBB3, GDF15, FHL1, NPY, DPP4, and GADD45A) and HIST2H2BE as a candidate one which are tightly correlated with the pathogenesis of PCa.

## Introduction

Worldwide prostate cancer (PCa) is recognized as the second most common diagnosed cancer and the fifth cause of cancer death among men globally, with an estimated 1.1 million global number of new cases diagnosed and 307,000 deaths in 2012 (1). The most important risk factors associated with PCa include age, family history, genetic factors, lifestyle, environmental influences, and diet. Rising incidence rates of PCa have been observed over the last few decades, largely due to screening and early detection procedures (2). Studies based on analyzing Nordic twin registries have demonstrated at least a 50% higher risk in monozygotic twins than dizygotic twins, suggesting that genetic rather than shared lifestyle factors are responsible for much of this familial aggregation (3, 4). Indeed, genetic factors have been estimated to be responsible for almost 42% of the risk (4). Cell survival and proliferation, acquired through deregulation of the cell cycle, are prerequisites for cancer (5, 6). The key molecules that regulate the cell cycle include cyclin-dependent kinases (CDKs) and their regulatory cyclin partners. Regulation of these complexes involves the control of cyclin production and its destruction through phosphorylation by specific kinases and other regulatory proteins (7, 8). Identification of potential and reliable biomarkers which are involved in the cell cycle and proliferation processes can be beneficial for early PCa detection, diagnosis and prognosis.

Over recent years, considerable researches have led to the development of several molecular and genetic assays that have provided a prospective direction for the development of prostate cancer biomarkers (9). High-throughput genomics technologies (e.g., gene expression microarrays) have been significantly changing biomedical research nowadays, which the expression patterns of thousands of genes can be monitored simultaneously (10). The emergence of novel high throughput technologies make the rapid identification of a single or groups of biomarkers possible (11, 12). Identification of differentially expressed genes (DEGs) often leads to the identification of novel cancer biomarkers followed by detecting upstream cancer causal genes by subsequent network analysis (13). In the current study, we downloaded two transcriptomic datasets from the Gene Expression Omnibus (GEO) in order to systematically detect DEGs. We constructed protein-protein interaction (PPI) network of the DEGs. Then, the gene ontology and Wikipathway of DEGs were done for the enrichment part. To get this goal, computational systems biology were applied to obtain a number of primary PCa DEGs from the results of transcriptomic analysis. Then, our findings may reveal the potential biomarkers in primary PCa detection and diagnosis.

## Methods

### Database Searching and recognizing pertinent experiments

From Gene Expression Omnibus (http://www.ncbi.nlm.nih.gov/geo/) database (GEO), we downloaded two PCa-related microarray datasets containing high quality transcriptomic samples concordance to the study design which compared the transcriptome of patients with the normal individuals. Searches were filtered for Homo sapiens and prostate cancer (PCa), benign and primary were the key words utilized in the search. Microarray raw data with accession numbers GSE6604, GSE6606 and GSE69223 were downloaded from this database. GSE69223 dataset (first dataset) which compared the transcriptome of 15 prospectively collected. It contained 15 benign PCa tumor biopsies and 15 normal samples from adjacent primary PCa tumors. GSE6604 included normal samples and GSE6606 contains primary PCa samples. So, we merged two datasets into a unified dataset (GSE6604GSE6606/Second dataset) consisting of 18 normal samples and 61 primary samples. All the samples ID are listed in supplementary file 1.

### Meta-Analysis and Identifying Differential Expressed Genes

R software was utilized to import and analyze data for each dataset separately. Two main Preprocessing steps involving background correction and probe summarization were performed using RMA method in Affy package (14). Absent probesets were also identified using “mas5calls” function in this package. For each probeset half of the samples had absent values, that probeset was regarded as absent and removed from the expression matrix. Besides, Outlier samples were identified and removed using PCA and hierarchical clustering approaches. Next, data normalization was done via quantile normalization method (15). Then, standard deviation (SD) for each gene was computed and SDs median was used as a cut off to remove low variant genes. NsFilter function in genefilter package was used for solving many to many problem (16) which is mapping multiple probesets to the same gene symbol (17). This function selects the probeset with the highest Interquartile range (IQR) between multiple probesets mapping to the same gene symbol. Limma package in R software was applied to perform linear models on the expression matrix data of each GEO dataset, and the DEGs between tumor and normal tissues were also screened by the limma package (18). |log2FC| >1, Benjamini Hochberg adjusted P-value < 0.05 (6) were considered statistically significant for the DEGs. Finally, common DEGs between two datasets were selected as the final DEGs.

### Network Construction

The STRING database was applied to identify potential interactions among the overlapping DEGs. The giant component of the weighted network was extracted from the whole network through using igraph package in R software (19). Next, the weighted adjacency matrix was transformed into a symmetric matrix to get modified into a new adjacency matrix using topological overlapping measure (TOM) function in WGCNA R package (20). To remain with a distance matrix, the new adjacency matrix was subtracted from one.

### Neighbourhood Ranking to the Core Genes

Using R software, a matrix of all shortest paths between all pairs of nodes in a weighted network was constructed using Dijkstra algorithm (21). Then, a distance score, Dj, for each node in the PPI network was calculated. Dj is the average of the shortest paths from all the non-core genes to reach the node j subtracted from the average of the shortest paths from the core genes to reach the node j normalized by the average of the all shortest paths to reach the node j from the whole network (22).

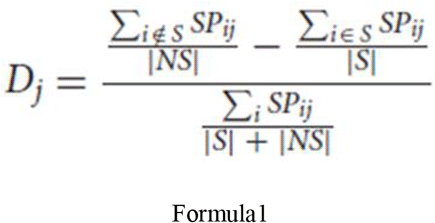

This scoring system implies how close each node is to the core nodes. Here, S is the set of core nodes and also NS is the set of non-core nodes. As a result, a score bigger than zero implicates that node j falls closer on average to the core nodes than it does on average to the rest of the network. Nodes with positive score were kept and the remained ones were deleted from the network. D scores were computed without imposing any threshold on edge weights.

### Enrichment Analysis

To explore the functions and pathways of DEGs, we carried out enrichment analysis. Enrichment analysis was performed using Enrichr online tool (23). Enriched terms for biological process were obtained from *GO* repository. For pathway enrichment analysis, DEGs were mapped into the *wikiPathways* signaling repository version 2019 for human. Enriched terms with the top score and the p-value less than 0.05 were selected.

## Results

### Data Preprocessing

Each dataset was imported into R separately. Almost 50% of probesets were regarded as absent and left out from the expression matrix to avoid technical errors to be engaged in downstream analysis. To be more precise in preprocessing step outlier sample detection was conducted using PCA and hierarchical clustering. Figure1A illustrates the PCA plot for the samples in first (GSE69223) dataset. The same plot was created for the samples in second (GSE6605GSE6606) study. Between two groups of samples, a number of them were away from their sets and should be considered as outliers. To be more specific, a hierarchical clustering method introduced by Oldham MC, et al (24) were used. Pearson correlation coefficients between samples was subtracted from one for measurement of the distances. Figure 1B depicts the dendrogram for the normal samples. Figure1C normal samples are plotted based on their Number-SD scores. To get this number for each sample, the average of whole distances was subtracted from distances average in all samples, then results of these subtractions were normalized (divided) by the standard deviation of distance averages (24). Samples with Number-SD less than negative two which were in a pretty long distance from their cluster set in the PCA plot, were regarded as outlier. Moreover, for every dataset, box plots were plotted and samples with extreme IQRs were removed. For instance, GSM1695600 was regarded as outlier based on the PCA and boxplot while GSM1695586 was considered as outlier based on Number-SD score. However, boxplot alone for some samples was sufficient to consider them as outliers regardless of PCA or Number-SD plots. Six outlier samples in first dataset and 31 outlier samples in second dataset were recognized. Supplementary file1 contains information about groups of samples and outliers.

**Figure 1:**
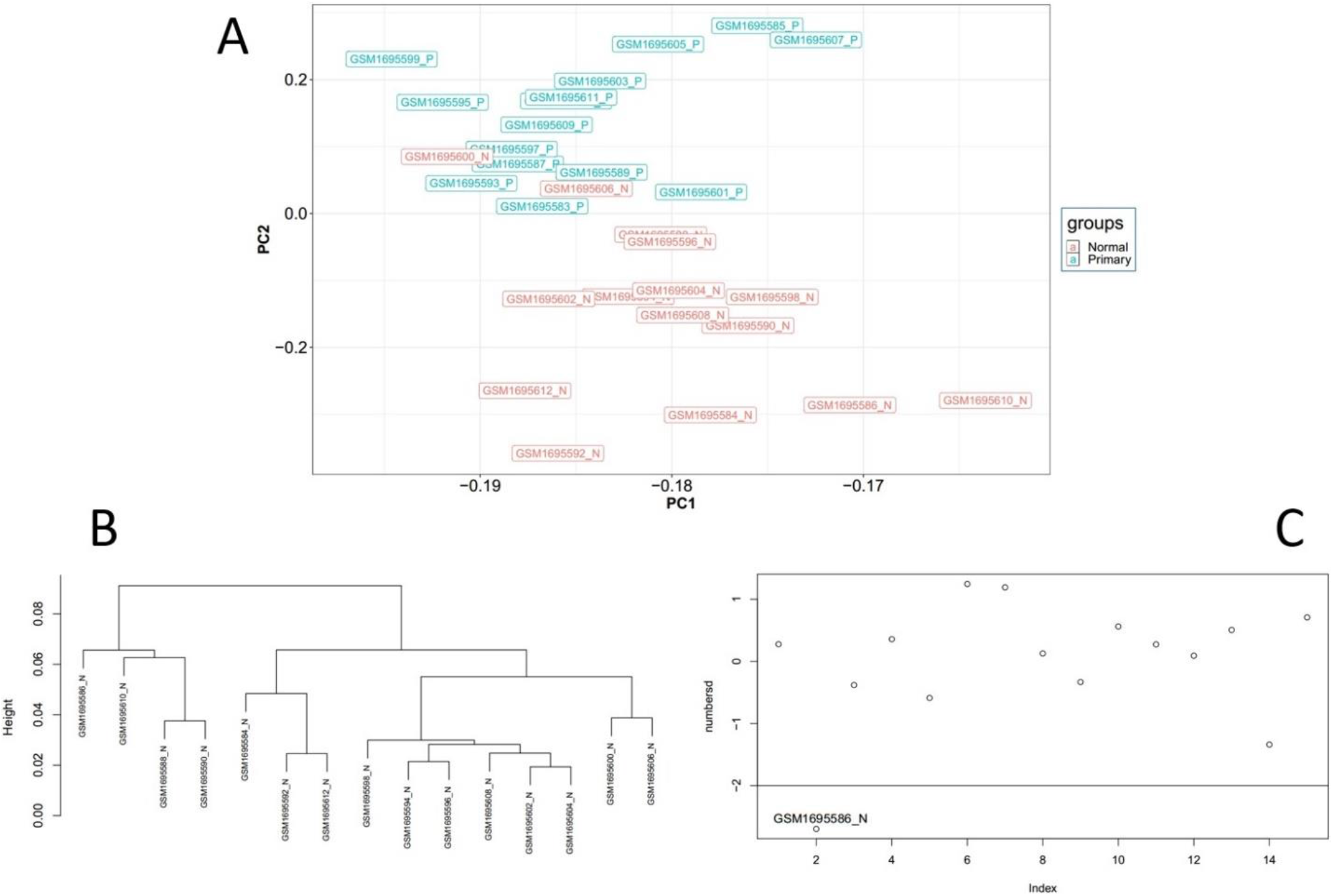
Illustration of outlier samples in first dataset. A, is the PCA plot, B is the dendrogram for the normal samples and C is the Number-SD plot for normal samples.

### Identifying Differentially Expressed Genes

All the samples in two groups including primary and normal were put into the calculation of DEGs. Genes with absolute log fold change (LogFC) larger than 1 and adjusted p-value less than 0.05 were regarded as DEGs. All 268 samples were put into the calculation of DEGs. Totally, 504 upregulated genes and 909 down-regulated genes were recognized in the first dataset as well as 154 overexpressed genes and 145 down-regulated genes in the second dataset. The final DEGs were selected from 60 common upregulated and 72 common down-regulated genes between datasets. DEGs with the absolute LogFC larger than 90% quantile of all DEGs (final DEGs) were considered as the core genes. This threshold was 1.919744 for up-regulated and −1.998201 for down-regulated genes in the first dataset and 3.22436 for up-regulated and −2.37239 for down-regulated genes in the second dataset. The core genes are shown in the scatter plots in Figure 2. Housekeeping genes are situated on the diagonal of the plot whilst up-regulated DEGs are above the diagonal and down-regulated DEGs are under the diagonal.

**Figure 2:**
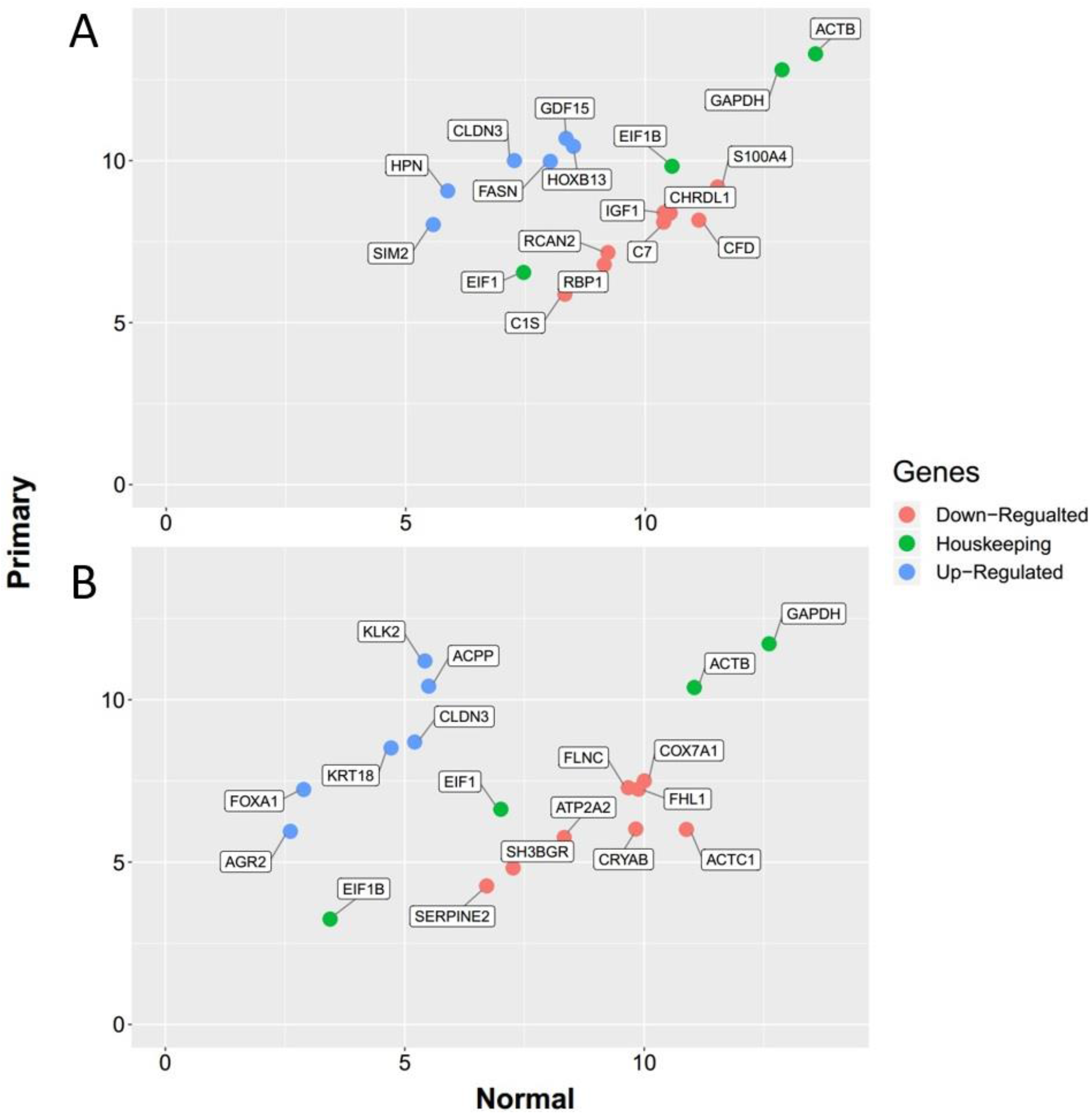
The quality Control plot. The housekeeping genes are near the diameter of plot presenting close average values between three groups.

### Undirected Protein-Protein Interaction Network

1575 unique DEGs with adjusted p-value < 0.05 and absolute log fold change > 1 were achieved from two sets of DEGs. Primary samples were compared to normal samples in both datasets. In the next step, 132 common DEGs between two sets of DEGs were retrieved. All DEGs for each dataset are presented in Supplementary file2. Common ones were employed to construct the Protein-Protein-Interaction (PPI) network. STRING database was used to generate the interactions based on seven filtrations namely, Neighborhood, Text mining, Experiments, Databases, Co-expression, Gene fusion, and Co-occurrence. STRING combined scores were used as the edge weights. This network involved 132 nodes and 248 edges. The giant component of this network with 97 nodes and 243 edges is presented in figure 3.

**Figure 3:**
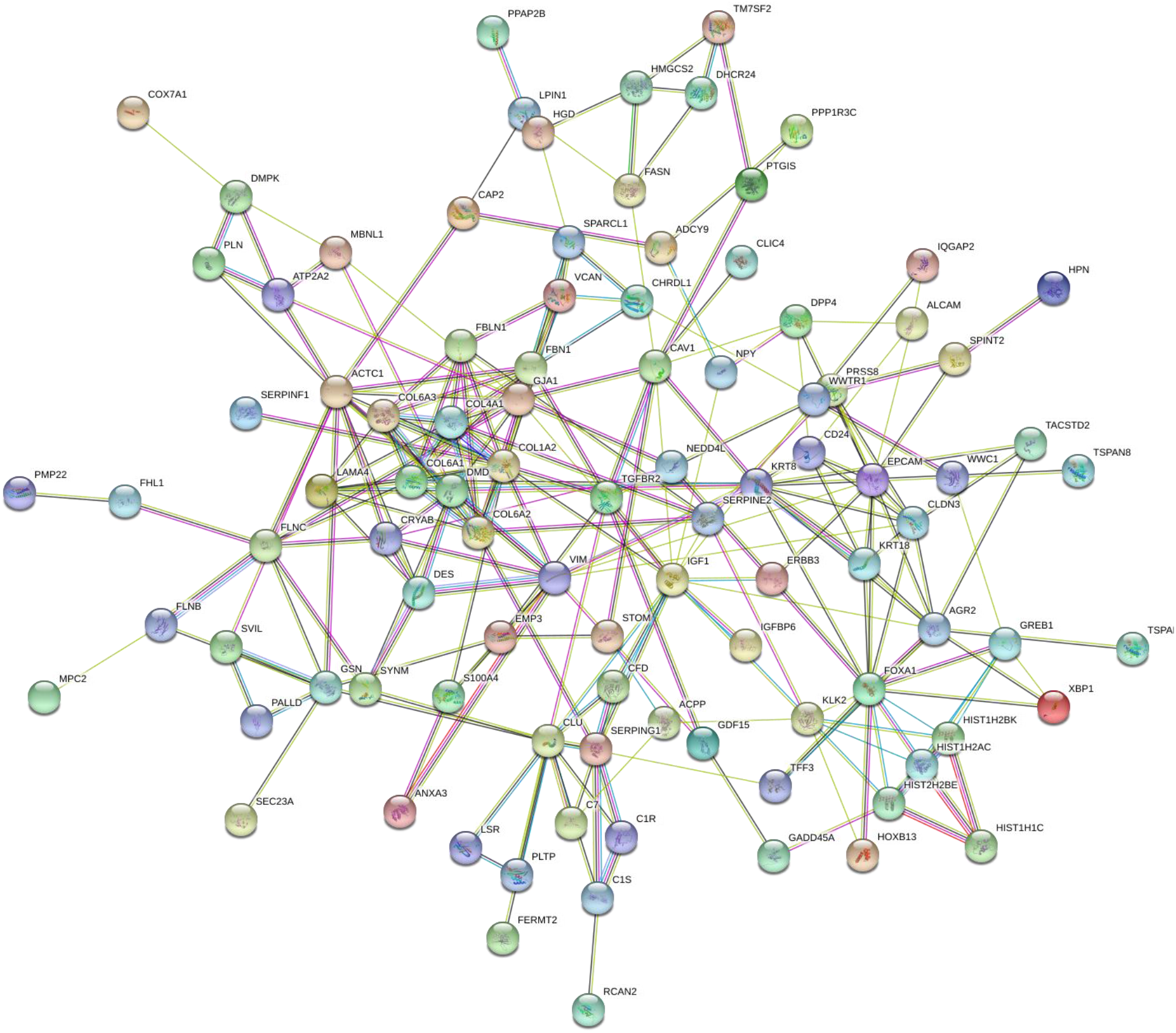
The whole network giant component. Labels are protein/gene symbols and edges with stronger evidence are thicker. This is a scale free network (25) which follows a power law distribution (most of the network nodes have a low degree while there are few nodes with high degree).

### Determination of Core Genes Neighborhood through Shortest Path-Based Scoring System

In this step, we used weights between nodes as the estimation of distances in the weighted adjacency matrix. Nodes with shorter distances from the core genes were selected and a smaller network was extracted from the main network. In order to choose core genes, 6 common DEGs in the first quartile of the most upregulated or downregulated genes in two datasets were considered. Computing the shortest path score for the non-core genes, led to a network with 24 nodes consisting of 6 core nodes and 22 neighbors. This two-component graph is called Core network shown in figure 4A. The expression states for these genes are illustrated in Figure 4B. In addition, two important centralities including degree and betweenness for Core network nodes are depicted in figure 4B. FOXA1 one of the core genes has the highest value in both centralities. ARG2 and EPCAM have pretty high degree and betweenness centralities. HIST2H2BE and DPP4 have relatively high betweenness value.

**Figure 4:**
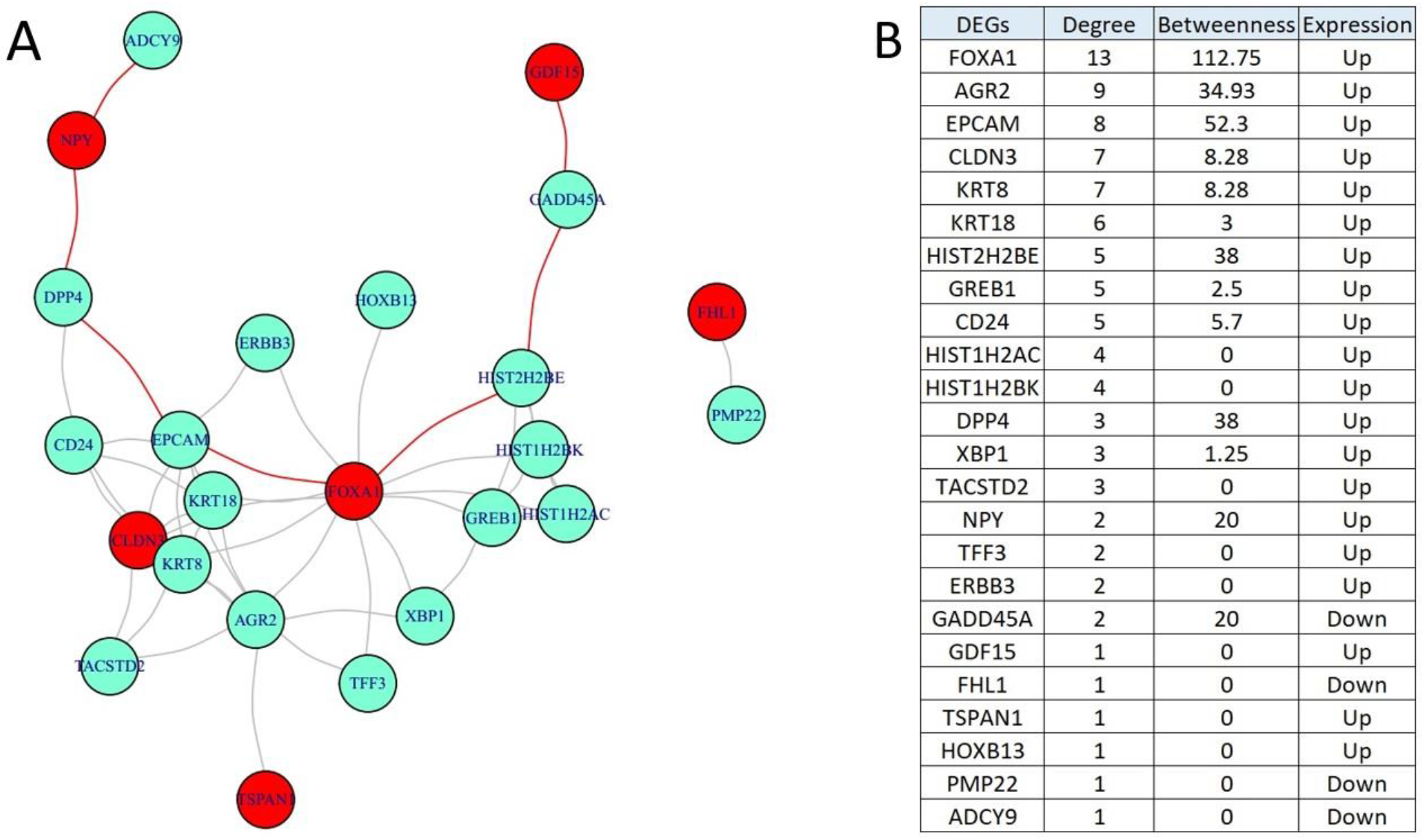
The Core network and centralities. This network presented in A contains two components and core genes are in red. B, Illustrates the Degree and Betweenness centralities for the nodes in the Core network. DEGs are sorted based on the highest Degree.

### Network Descriptive

The Core network diameter is seven containing GDF15, GADD45A, HIST2H2BE, FOXA1, EPCAM, DPP4, NPY, and ADCY9. The giant component descriptive is explained in the following. Transitivity was around 52%, edge density was about 17.4% and the mean distance is 2.35. Other centralities for the nodes and the average distances between each node and the other nodes are provided in Supplementary file3.

### Network Clustering and Enrichment Analysis

Giant component (GC) of Core network were clustered using fast greedy algorithm in R. It is illustrated in Figure 5. Gene Set Enrichment Analysis (GSEA) was performed on each cluster separately. Gene sets in clusters 1 to 3 were given to *Enrichr* online tool. Tables 2A and 2B depict the enrichment analysis for the core network giant component. Enriched terms are sorted based on the highest p-value.

**Figure 5:**
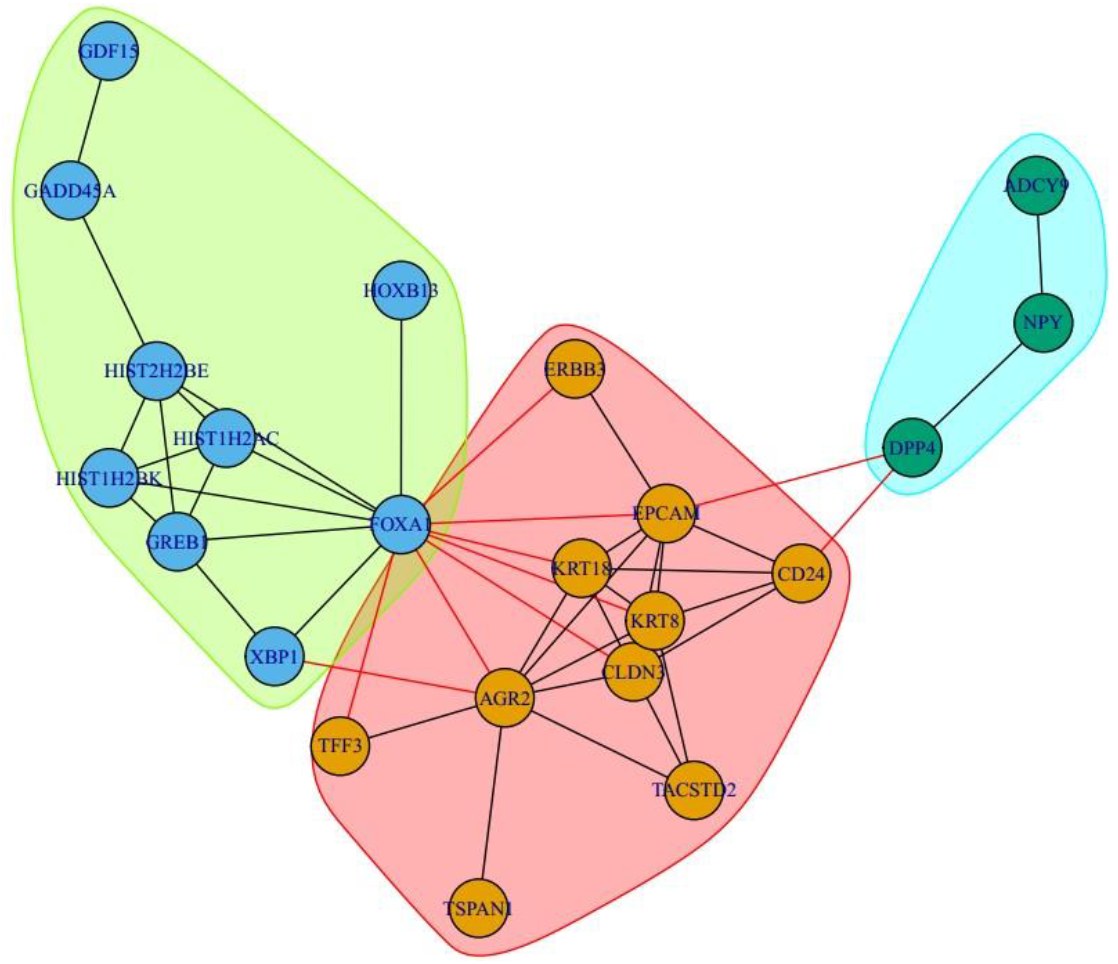
Clustering results for GC of the Core network. Nodes in each cluster are situated in the same color aura. First cluster is in red. Second cluster is in green and third cluster with three nodes is in blue.

Apoptosis-related network due to altered Notch3 in ovarian cancer, Signaling Pathways in Glioblastoma, and ERBB3 Signaling Pathway are enriched in Wikipathways containing ERBB3, KRT18, and KRT8 genes. ERBB3 is involved in all 3 pathways. Engagement of this gene in pathways-related cancer (WP2864, WP2261, and WP673) and involvement in the biological processes including regulation of cell motility (GO:2000145), regulation of MAPK cascade (GO:0043408), and regulation of signal transduction (GO:0009966) proposes the potential importance of this gene in different cancers. In 2B, CD24 gene has been shown to monopoliz the most GO terms to its self, then the highest number of terms is related to ERBB3 and AGR2. The enrichment analysis for the core network components of 2 and 3 are presented in Supplementary file 4.

In component 2, two genes enriched in Wikipathays. One of these genes, GADD45A, is involved in 12 pathways whose involvement in all of them demonstrates the vital role of this gene in pathogenesis and progression of cancer (TP53 Network WP1742, Imatinib and Chronic Myeloid Leukemia WP3640, ATM Signaling Pathway WP2516, OEndometrial cancer WP4155, G1 to S cell cycle control WP45, Chromosomal and microsatellite instability in colorectal cancer WP4216, and …). Moreover, this gene is implicated in regulation of apoptotic process (GO:0042981) and activation of MAPKKK activity (GO:0000185) in aspect of biological process in GO. HIST2H2BE and HIST1H2BK genes have monopolized the most GO terms to its self. After them, XBP1 has the highest number of terms. In component 3, ADCY9 has enriched for the most pathways and possesses the most GO terms.

**Table 1:**
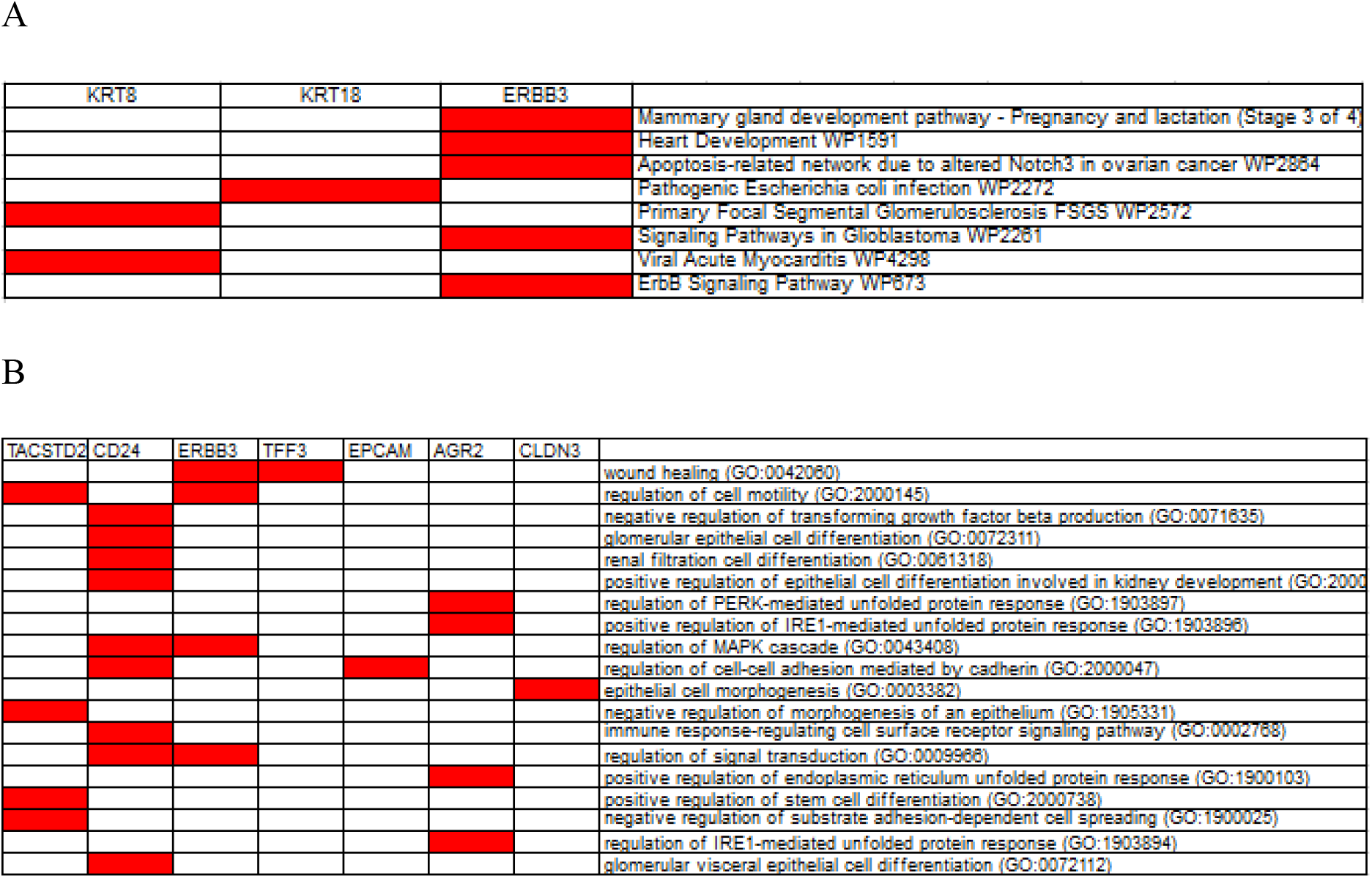
Enrichment terms for the giant component. 2A illustrates the enriched pathways from WikiPathways signaling datasets. 2B shows the enriched terms for biological process from GO repository. Red cells reveal the involvement of the genes in the enriched elements.

## Discussion

Integrated bioinformatics analysis mainly focusing on differentially expressed molecule screen has been extensively applied to identify potential biomarkers associated with the diagnosis, treatment, and prognosis of cancers. With two microarray datasets from the GEO database, we identified DEGs in primary PCa and normal tissues. Cell cycle related biological processes are highlighted as the significant items for primary PCa in this study. The association of all of these genes with different cancers has already been confirmed.

FOXA1 one of the core genes has the highest value in both centralities (degree and betweenness) and exhibited upregulation in PvsN analysis. FOXA1 is characterized as a ‘‘pioneer factor’’ having the special property of being able to directly bind DNA (26). Increased expression of FOXA1 has also been observed in colon, lung, thyroid, esophageal and prostate cancer (27). FOXA1 not only plays a role in androgen receptor (AR) signaling, but also regulates the expression of genes engaged in cell cycle regulation in prostate cancer (28–30). Imamura et all. identified IGFBP-3 as a down-stream target of FOXA1 (28). Increased IGFBP-3 expression level following reduction of FOXA1 significantly inhibited proliferation of PCs most likely via the phosphorylation inhibition of signaling mediators in the IGF-1 signaling pathway including MAPK and Akt, and mediated cell cycle arrest through p21 and p27 in PCa cells (28). This gene has been identified as not only an attractive therapeutic target but could potentially function as a biomarker in previous investigations (31).

AGR2, anterior gradient 2, is a member of protein disulfide isomerase (PDI) family resides in the endoplasmic reticulum (ER) (32). AGR2 expression has been found in prostate: primary and metastatic, pancreatic, breast, lung, gastrointestinal and oral solid tumors (33, 34). In PvsN analysis, overexprission of this gene has been observed and also had pretty high degree and betweenness centralities. The reduced level of AGR2 expression by using ARG2-specific siRNA led to cell cycle arrest at G 0 /G 1 phase, providing several pieces of evidence that AGR2 is a kind of cell cycle modulator (35). AGR2-silenced cells display significant reduced cell proliferation and invasion of pancreatic cancer cells (36). Arrest in G 0 /G 1 phase of cell cycle due to transient knockdown of AGR2 is correlated with senescence response in malignant prostatic cells (35). Cellular senescence is characterized as a key mechanism of tumor suppression (37). AGR2 has been suggested as a potential biomarker for the diagnosis of some cancers including pituitary adenomas, prostate, human lung adenocarcinoma, and breast (38–41)

Epithelial cell adhesion molecule (EpCAM), also known as CD326, is a transmembrane glycoprotein acts various roles including cell signalling, migration, proliferation and differentiation besides cell-cell adhesion (42). EpCAM signaling is activated upon regulated intramembrane proteolysis (RIP) and subsequent release of the intracellular domain EpICD, which associates with components of the Wnt pathway (FHL2, *β*-catenin, Lef-1), translocates to the nucleus, and functions as transcription regulator. EpCAM induced genes, such as *c-myc, cyclin D1, mmp7* (43–46), and regulators of the cell cycle and proliferation machinery, in general (47). Overexpression of EpCAM has been found to be associated with enhanced transcription and translation of the proto-oncogene *c-myc* (43). EpCAM was then found to be expressed at a high level and frequency not only on primary PCa (48) and colon cancer tissues but on most human adenocarcinomas (49) as well as on squamous cell carcinomas (50). In current study, Increased EpCAM expression has been observed and had been identified as a biomarker for hepatocellular carcinoma and prostate cancer (51, 52)

NPY is a 36 amino-acid sympathetic neurotransmitter which is abundant in the brain (53). NPY has been revealed to be engaged in mitogenic pathways and stimulate cellular proliferation through the Y1-R, its specific G protein-coupled receptor (54). The Y1-R activation is generally associated with decrease of cAMP accumulation, increase of intracellular free calcium concentration ([Ca^2+^]_i_), and modulation of the MAPK pathway via several signaling molecules, including the protein kinase C (PKC) (54, 55). Recent researches have shown that NPY can promote cell proliferation of endothelial and increase tumor vascularization through VEGF pathway in some solid tumor cell types (56, 57). The mRNA expression level of *NPY* is significantly higher in non-aggressive PCa cells than that in aggressive PCa (58). Elevated level of NPY expression has been found in current study and also some studies have found NPY receptors to be overexpressed in neuroblastoma and breast cancers, and emerged as promising targets in cancer diagnosis and therapy (59–61)

*CLDN3* belongs to a family of proteins engaged in the formation and function of tight junctions (TJs) (62). Hewitt *et al.* reported CLDN3, CLDN4, and CLDN7 to be overexpressed in the pancreas, stomach, colon, bladder, ovary, breast, uterus, and in both primary and metastatic prostate cancer and also it has been overexpressed in PvsN analysis (63, 64). Overexpression of *CLDN3* is regulated by the EGF-activated downstream pathway, ERK1/2 and PI3K-Akt signaling pathways (65). More importantly, the EGF-induced overexpression of *CLDN3* in adenomacarcinoma (ADC) cells reinforces the cell proliferation (65). *In* another *study CLDN3* was suggested to be a potential molecular target for future application in the treatment of ADC (66). It is interesting that almost all of these cell cycle regulatory proteins can be modulated by PI3K signaling (67). ERBB3 can interact with many genetic and signaling pathways, including the PI3K/AKT pathway (68, 69). In recent study, G2/M arrest occurred in both MKN45 and SGC7901 by silencing ERBB3. Since G2/M transition is mediated by CDK1, by decreasing cyclinB1, one of its most important regulators, activity of CDK1 was also decreased. Inactivation of Akt by expression of its dominant negative mutant form inhibited cell proliferation by arresting the cells at G2/M phase through decreasing cyclinB1 (70). Overexpression of ERBB3 was reported in different types of cancer including bladder, prostate and breast cancer as well as current study (71, 72). In enrichment analysis, ERBB3 is involved in three pathway-related-cancer (WP2864, WP2261, and WP673). This gene is a biomarker for some cancers including oropharyngeal squamous cell carcinoma and pancreatic cancer (73, 74).

*GDF15,* which encodes growth/differentiation factor-15, is a cytokine and member of the transforming growth factor beta (TGFβ) family (75). GDF15 expression is downregulated in metastatic PCa in comparison with the primary tumors, levels of GDF15 RNA and protein are higher in primary prostate tumors (76). In agreement with previous results, elevated GDF15 expression has been found in our PvsN analysis. In prostate cancer, maspin inhibits tumor growth, reduces bone metastasis, and decreases angiogenesis in vitro and in vivo (77–79). Based on the result of an investigation, overexpression of GDF15 blocked maspin gene expression. This finding might explain how overexpression of GDF15 leads to an increased proliferation and invasiveness of prostate carcinoma cells by GDF15 (80). Elevated expression of GDF15 has been found in the samples of patients with breast and colorectal carcinomas (81). In some studies, GDF15 has been identified as diagnostic and prognostic biomarker for prostate, colorectal and ovarian cancer (82–84)

FHL1 is significantly down-regulated expressed and plays important roles in the regulation of development and progression in several types of human tumors including prostatic carcinoma, bladder cancer (BC), breast carcinoma, and prostate and the same result for the current research has been obtained (85–89). FHL1 impacts the progression of cancers by cell cycle regulation of tumors. FHL1 increases the expression of p21 (a CDK inhibitor) through regulation of gene transcription. In addition, p21 plays a role in the maintenance of G2/M-phase arrest, and regulation of cell cycle arrest at the G1/S transition. FHL1 induces the upregulation of p21 and inhibits cyclins and their associated CDKs (mainly cyclin D1), which are the central machinery governing cell cycle progression (90). FHL1 plays important roles not only in activation of the tumor suppressor gene *p21,* but also in repression of the oncogene *c-myc* through interacting with Smad2, Smad3 and Smad4 that are important regulators of cancer development and progression in a casein kinase 1ô-dependent manner (91). It has been emerged as a biomarker for progression of cutaneous squamous cell carcinoma (92).

Growth arrest and DNA-damage-inducible protein 45 alpha (GADD45A) is a downstream target gene of p53 and BRCA1 (breast cancer susceptibility gene 1) (93). GADD45 family involve in cell cycle progression, cell survival and apoptosis (94, 95). By interaction with Cdc2 kinase, Gadd45a can dissociate Cdc2/cyclinB1 complex and mediate G2/M cell cycle arrest (96, 97). In addition, GADD45a was reported to suppress the tumor angiogenesis through downregulating of VEGFa via blocking the mTOR/STAT3 pathway (98). Downregulation of GADD45 expression can be possible by the activation of several oncogenic signaling molecules, such as c-Myc, nFκB or Akt (99). GADD45A protein levels were increased significantly in breast cancer (100) compared with normal tissues and also highly expressed in human pancreatic cancer (101), whereas downregulated in bladder cancer tissues (102). In PvsN, decreased expression level of GADD45A was found. Moreover, it has been suggested as a biomarker for PCa (103). In component 2, GADD45A, was enriched in Wikipathays and involved in 12 pathways associated with cancer and its engagement in all of them demonstrates the vital role of this gene in pathogenesis and progression of cancer.

Dipeptidyl peptidase IV (DPPIV), a membrane glycoprotein, controls the activities of mitogenic growth factors and neuropeptides. DPPIV is involved in various biological processes, including cell differentiation, adhesion, immunomodulation, and apoptosis, functions that are critical for the neoplastic transformation regulation (104–106). In prostate cancer cells, overexpression of DPIV is found in benign glands, whereas little or no DPIV is detected in prostate tumor metastases (107). In aspect of expression, the same result has been found in the current study. DPIV up-regulation in prostatic tumors influences tumor growth through two possible mechanisms based on its potential in peptidase activity, that leads to increase cleavage of a propeptide to its active form could stimulate cell proliferation or alternatively DPIV could inactivate a peptide that inhibits cell proliferation (108). DPP4 could be a potential biomarker and target for cancer therapy (109).

In humans, histone H2B is coded by twenty-three different genes, none of which contain introns. All of these genes are located in histone cluster 1 on chromosome 6 and cluster 2 and cluster 3 on chromosome 1 (110). This gene was upregulated in our analysis which is connected to other histone core genes namely HIST1H2BK and HIST1H2AC in Figure 5. This gene also is connected to GADD45A a growth arrest gene. A correlation between reduced expression of Gadd45 and increased resistance to topoisomerase I and topoisomerase II inhibitors in a variety of human cell lines was found. Gadd45 could potentially mediate this effect by destabilizing histone-DNA interactions since it was found to interact directly with the four core histones (111). Gadd45 expression was down regulated in our analysis. This downregulation might associated with the HIST2H2BE upreglution but the mechanism behind this process is not documented yet and this needs a further investigation. However, more investigation needs to be conducted on relation of this gene since no study has connected HIST2H2BE to any kind of cancer so far. This gene has also a relatively high betweenness centrality and HIST2H2BE and HIST1H2BK genes have monopolized the most GO terms to its self which are related to the immune response and DNA assembly. Moreover, it can be proposed as a possible diagnosis and prognosis biomarker for PCa.

In conclusion, with the employment of multiple gene expression profile datasets and integrated bioinformatics analysis, we have demonstrated 10 potential biomarkers which have been confirmed by previous investigations. In addition, HIST2H2BE has been identified as a possible biomarker for the pathogenesis of PCa. The results of this study can shed more light on common molecular mechanisms adopted by PCa.

